# Systematic promoter perturbation reveals limitations of AlphaGenome gene expression predictions

**DOI:** 10.64898/2026.11.06.764851

**Authors:** Varun K A Sreenivasan, Ekin Deniz Aksu, Malte Spielmann

## Abstract

Sequence-to-function (S2F) models promise to facilitate the interpretation of noncoding variants by predicting their effects on gene regulation directly from genomic sequence. However, whether these models recapitulate fundamental principles of transcriptional biology remains unclear. Here, we tested whether AlphaGenome recognizes the expected consequences of promoter disruption on predicted gene expression. We systematically perturbed annotated promoter elements *in silico* for 2,111 promoter–gene pairs using dinucleotide shuffling, GC-adjusted random base substitution, and N-masking. We then compared AlphaGenome-predicted CAGE and RNA-sequencing signals across 20 tissues for reference and promoter-disrupted sequences. While CAGE and RNA-sequencing predictions were strongly correlated for reference sequences, promoter disruption markedly reduced predicted CAGE activity, with N masking producing reductions of approximately three orders of magnitude. By contrast, the predicted RNA-sequencing signals were substantially less affected, resulting in an uncoupling of two molecular readouts of gene expression. Exon-level analyses showed that AlphaGenome predicted reduced RNA-sequencing signal when exonic sequences are within the disrupted region, but frequently failed to propagate this effect to downstream exons. This divergence between the CAGE and RNA-sequencing readouts was apparent from the second exon onward and was more pronounced in longer genes. Although alternative transcription initiation and shared downstream exons may explain individual cases, it is unlikely to account for the systematic discordance between CAGE and RNA-sequencing predictions. These findings indicate that AlphaGenome can recognize the local loss of promoter activity without consistently capturing its consequences across the corresponding transcript. More broadly, our results show that predictions from individual S2F output tracks do not necessarily reflect a coherent model of transcription. Systematic biological benchmarking and experimental validation therefore remain essential before such predictions can be used to prioritize or classify noncoding variants in research or diagnostic settings.

## Main text

Interpreting the molecular and phenotypic consequences of genomic variation remains a central challenge in human genetics, particularly for variants outside protein-coding regions^1^. Experimental assays in cellular and animal models provide essential evidence, but cannot be readily scaled to the vast number of variants identified by genome sequencing. Computational approaches are therefore increasingly used to predict molecular effects and prioritize candidate disease-causing variants^2^. Earlier methods typically considered individual features, including predicted effects on protein function^3^ (e.g. PolyPhen^4^), and evolutionary conservation^5,6^. Subsequent machine-learning approaches integrated multiple annotations but were initially developed and most extensively validated for protein-coding variation^7^. More recently, sequence-to-function (S2F) models trained on large collections of functional genomic data have promised to extend variant-effect prediction across the noncoding genome. Models such as Enformer^8^, Borzoi^9^, ChromBPnet^10^, DeepSTARR^11^, Orca^12^, and AlphaGenome^13^ predict diverse molecular readouts—including gene expression, chromatin accessibility, and three-dimensional genome organization—directly from genomic sequence and, in some cases, across different cellular contexts. Analysis indicates that S2F models can classify causal single-nucleotide variants (SNVs), make reasonably accurate predictions of expression quantitative trait loci (eQTL) effect direction, and recognise regulatory grammar such as transcription factor binding motifs and topologically associating domain boundaries^8,9,13–18^. Hence, these models are increasingly being applied to prioritize rare and personal variants and to design synthetic regulatory sequences, reflecting the expectation that they capture fundamental aspects of the *cis*-regulatory grammar of the human genome. The recently released AlphaGenome Atlas^19^ further expands the accessibility of such predictions by providing estimated effects for every possible single-nucleotide variant and numerous observed indels across the human genome.

The ability to predict regulatory consequences directly from sequence could substantially reduce the experimental burden of interpreting noncoding variants and make S2F models highly attractive for research and clinical diagnostics. Their safe application, however, requires a clear understanding of where their predictions are biologically reliable and where they fail— particularly because model outputs remain difficult to interpret and generally lack calibrated measures of confidence. Several independent studies have benchmarked S2F variant-effect predictions beyond the evaluations presented with the original models^20–22^. They generally compare predicted effects with experimental or population-based reference datasets, including massively parallel reporter assays, fine-mapped eQTLs, saturation mutagenesis experiments, and CRISPRi-derived enhancer–gene maps^23–25^. While S2F models have shown to perform remarkably well in specific benchmarks, they tend to underestimate the magnitude of variant effects on gene expression^26^, with performance often declining as the distance between the perturbed *cis*-regulatory element and its target gene increases—for example, for distal enhancers compared with proximal regulatory sequences^9,13,18,20,21^. However, such benchmarking is constrained by the limited scale and context specificity of available experimental datasets, as well as by batch effects and the statistical assumptions used to define reference variant effects.

Therefore, we considered a complementary strategy that does not depend exclusively on individual variant-effect datasets. Instead, we asked whether application of AlphaGenome at promoter sequences would reproduce established features of transcriptional biology. Promoters provide an especially informative test case: they are among the best-characterized regulatory elements^27^, and their sequence architecture, transcriptional orientation, and relationship to gene expression are supported by extensive experimental evidence^28–32^. Assessing whether AlphaGenome recapitulates these established principles of promoter biology therefore provides a test of internal model consistency and whether its predictions can be interpreted as biologically meaningful evidence for noncoding variant effects.

Here, we analyse the predicted effect of severe promoter disruption on gene expression. This *in silico* experiment can help analyse the congruence between the model and RNA transcription in two aspects, namely (1) whether the two measures of gene expression, namely CAGE and RNA-seq, are predicted to be similarly affected upon promoter disruption, *i*.*e*. whether their correlation is maintained. This is relevant because it is reported that predicted variant effect is dependent on which functional track is considered^22^ and (2) to what extent the model recognises the loss of promoter region, while predicting the expression at distal downstream exons in the RNA-seq tracks. The latter indicates the model’s understanding of 5’ to 3’ directionality and continuity during gene transcription. We used AlphaGenome as a representative state-of-the-art model. Given a DNA sequence of up to 1Mb in length, AlphaGenome can predict ∼7k functional tracks that include gene expression, splicing patterns, or 3D genome contacts across multiple cell/tissue types in humans and mice.

We limited our analysis to a subset of human protein-coding genes, chosen at random. For each gene, we defined the promoter regions that were to be disrupted in the following manner. First, Eukaryotic Promoter Database (EPD v006) core promoters^27^ (with gene associations) were used as the basis for identifying the core promoters, with some genes annotated with as many as six sites. Next, to limit transcriptional activity from neighboring *cis*-regulatory sequences, all ENCODE candidate *cis*-regulatory elements with promoter-like signatures (PLS) and proximal enhancer-like signatures (pELS) that were in the vicinity (< 1kb and < 2kb, respectively) of the EPD core promoter were also defined as promoter elements of that gene. We also defined all robust FANTOM5 CAGE DPI peaks with matching gene annotations^33^ as promoter elements. Since many of these annotations were overlapping, we reduced the individual elements to overlapping promoter elements, whose sequences were then disrupted by various sequence perturbations detailed below (Figure 1b). Most genes had ∼10 non-contiguous promoter elements that were disrupted. All genes for which the region spanned by the promoter elements exceeded 10 kb and those present in duplicates at pseudoautosomal regions were not further considered for the analysis, resulting in a total of 2108 promoter-gene sets that were analysed. We also included three additional genes (*APOL4, RGS11*, and *ZNF155)* highlighted in the original publication in the context of RNA-seq specific variant effect prediction^13^.

**Figure 1.**
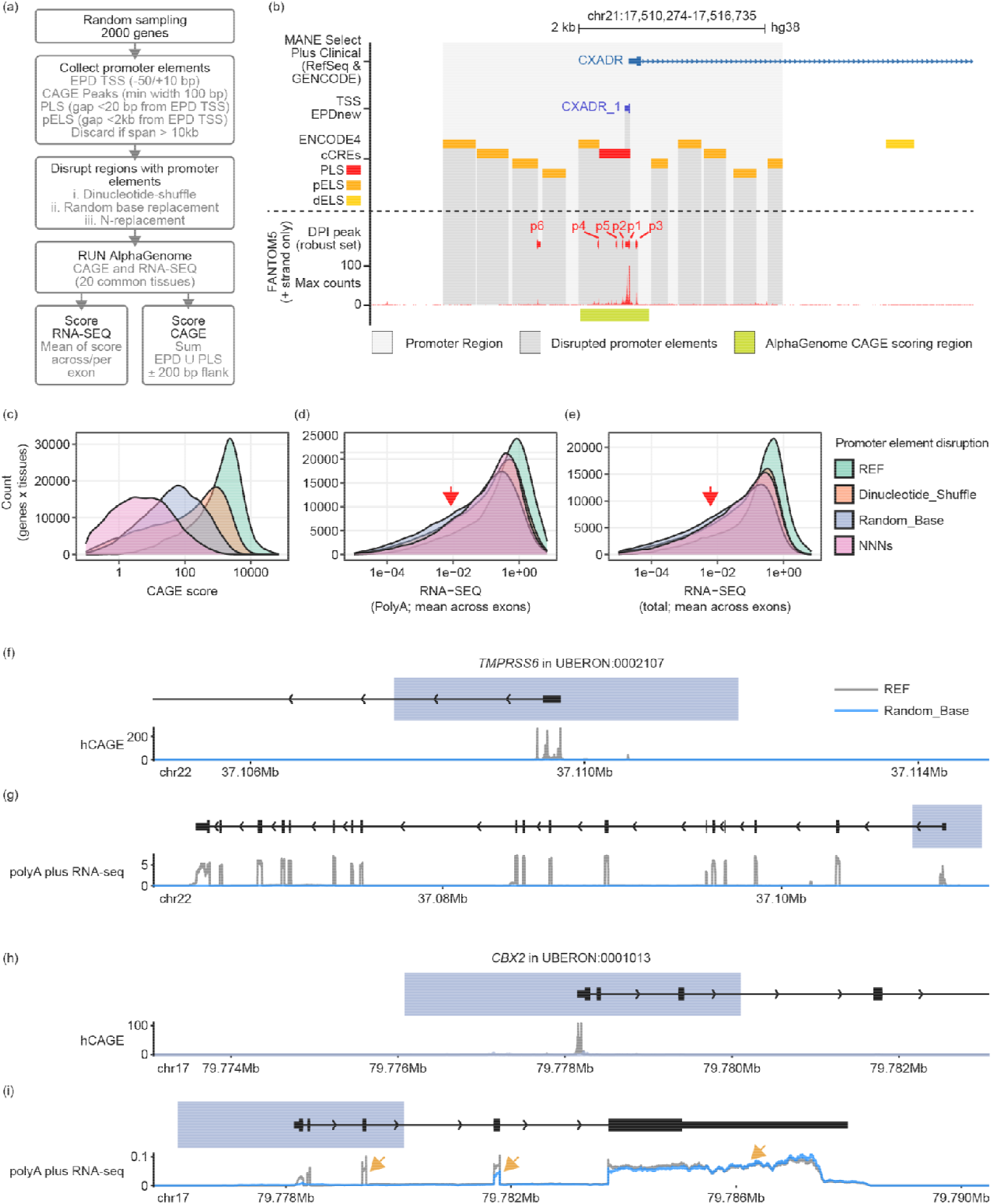
(a) A schematic of the approach used for evaluating the sensitivity of AlphaGenome to promoter sequence disruptions. (b) Promoter elements of genes to be disrupted (dark grey) were defined based on a set of annotations including EPD, ENCODE PLS and pELS, and FANTOM5 CAGE peaks. CAGE tracks predicted by AlphaGenome were scored by summing the scores in the CAGE scoring region (green), defined as range spanned by EPD and ENCODE PLS with an additional 200 bp flank on either ends. (c-e) Distribution of (c) CAGE, (d) polyA plus RNA-seq, and (e) total RNA-seq scores for the reference or promoter-disrupted sequences across all tested promoters and tissues. Note, total RNA-seq scores were not available in all 20 tissues. Red arrows highlight the more pronounced tail upon promoter disruption. (d-e) Representative examples of genes that (d) responded or (e) not to GC-adjusted random base promoter disruption in the predicted RNA-seq track. The UBERON id indicates the tissue for which this track was predicted.

We then disrupted these promoter elements in three different ways. First, we used dinucleotide shuffling, which is a common approach to disrupt regulatory motifs by randomizing the sequence while maintaining the ratio of dinucleotides^34^. However, many core promoters, especially of housekeeping genes, are demarcated by CpG islands with high GC content^27^, where dinucleotide shuffling may be ineffective in disrupting promoter activity. For example, in the regions that we identified as promoter elements, the GC fraction reached ∼55% in comparison to ∼42% in the undisrupted sequences in between. Therefore, to achieve a stronger promoter disruption we applied a GC-adjusted random base substitution approach, where the bases in promoter elements were replaced with random bases, but with the GC content constrained at ∼42%. Finally, we also analysed the effect of replacing the promoter elements with Ns, which effectively removes all promoter-like sequences. We reasoned that this approach is likely to be the most consequential to promoter activity disruption, given that Ns are numerically represented as zero vectors in the one-hot sequence encoding and are used in the model training as padding for short input sequences^13^. No insertions or deletions were introduced in either of the three disruption approaches, thus maintaining the genomic coordinates for scoring purposes.

We predicted CAGE and RNA-seq tracks for the reference and the promoter-disrupted sequences at each of the 2111 promoters using AlphaGenome, running locally on an NVIDIA A100 GPU (CUDA version 12.8). We have also previously verified that track predictions obtained with local installation were comparable to that obtained using the AlphaGenome cloud service, and we could reproduce the results presented in Figure 4 of original publication using the local installation (*APOL4* variant, data not shown). Since one of our goals is to compare the predictions across CAGE and RNA-seq, we limited the analysis to 20 tissues for which both CAGE and polyA plus RNA-seq tracks were available. A single score was calculated for each of the predicted CAGE and RNA-seq tracks similarly to the recommendation by the authors. Specifically, the CAGE score per tested promoter sequence was defined as the sum of predicted signals in the CAGE scoring region, which spanned the regions containing all EPD and ENCODE PLS sites and an additional 200 bp flank on either ends (Figure 1b). polyA plus and total RNA-seq scores for the tested promoter sequence were defined as the mean predicted value across all the exons of the RefSeq + MANE Select transcript associated with that promoter.

First, we verified that the promoter disruption resulted in reduced CAGE scores (Figure 1c, left). As expected, the GC-adjusted random base disruption yielded a larger reduction in CAGE scores compared to dinucleotide shuffling. However, the N-replacement approach was most effective in disrupting the predicted CAGE activity, reducing the score three orders of magnitude. In comparison the distributions of both in polyA plus and total RNA-seq scores were much less affected by the three promoter disruptions (Figure 1d,e). We found this divergence in responses between RNA-seq and CAGE surprising, as they are both quantitative measures of gene expression and are strongly correlated when using reference sequences (Figure 2c). Interestingly, the distributions in the case of disrupted promoters exhibited a more pronounced tail to the left (red arrows), indicating the presence of “responding” and “non-responding” genes, best visualised in the case of examples (Figure 1f-i). Disrupting the promoter of *TMPRSS6* (responding gene) with the GC-adjusted random base substitution strategy resulted in the severe reduction of both predicted CAGE and RNA-seq signals, whereas in the case of *CBX2* (non-responding gene) AlphaGenome predicted residual RNA-seq signal at downstream exons, despite the CAGE signal being visibly abolished. Furthermore, the residual RNA-seq signal increases with increasing distance from the promoter: exons within the disruption region have no residual expression, the first exon downstream of the disruption is partially affected, while the last exon is unaffected. This contradicts the biologically-founded expectation of correlated CAGE and RNA-seq signals. Indeed, it has been reported that predicting personalized gene expression is inherently challenging for S2F models, partly since gene expression is dependent on distal *cis*-regulatory elements. In comparison, prediction of a more localized signal such as chromatin accessibility is easier learnt by the models^22^. We wondered if this might explain why CAGE scores, which in our case were measured at the same locations where the sequences were disrupted, respond more strongly than along the exons which might be up to several hundreds of kilobases away.

**Figure 2.**
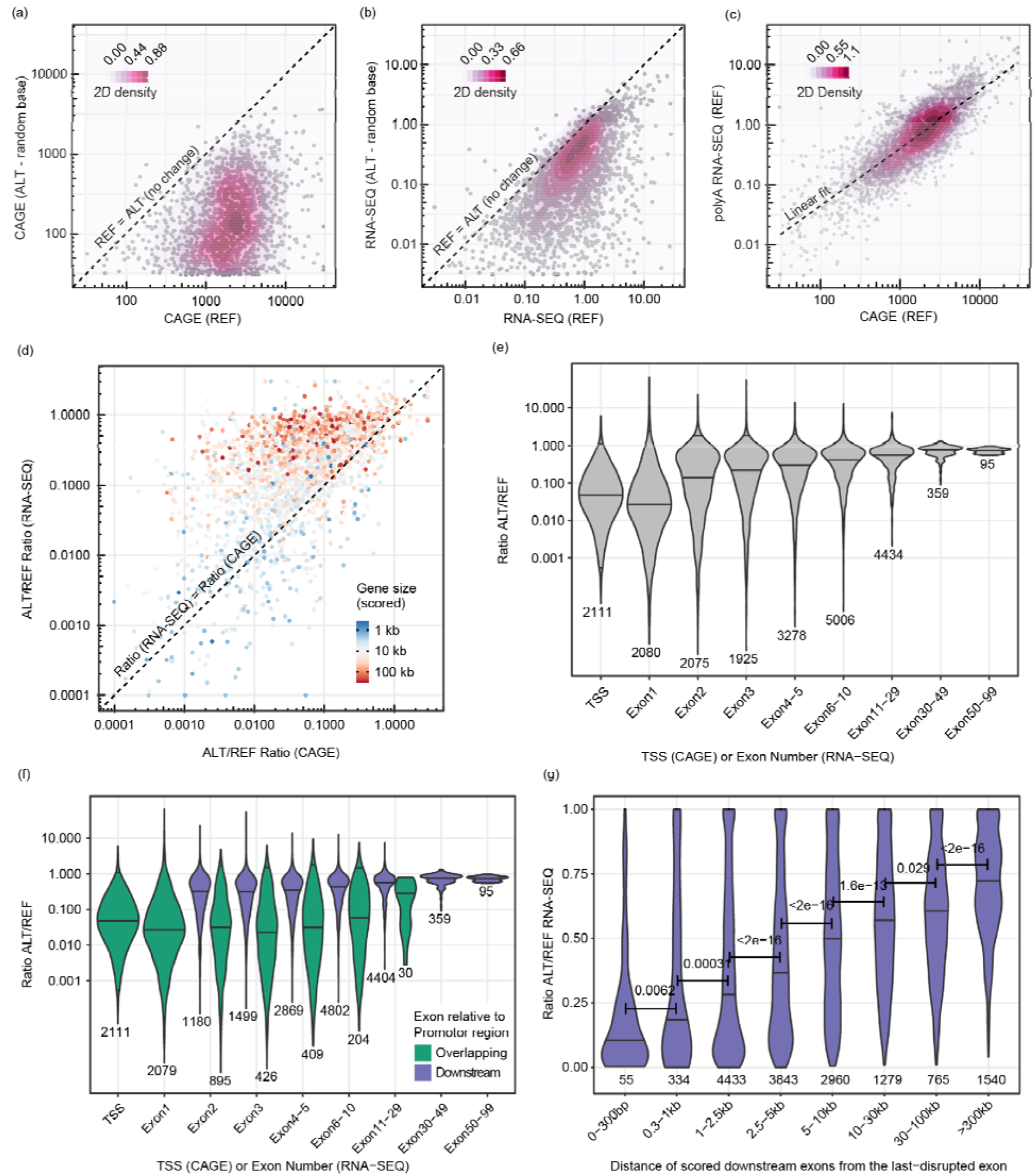
Analysis of the CAGE and polyA plus RNA-seq scores calculated from the respective tracks predicted by AlphaGenome for the human genome reference sequence (REF) and after GC-adjusted random base substitution (ALT). (a, b) Comparison of scores in REF and the ALT sequences, where points below the dotted diagonal indicate reduced (a) CAGE or (b) RNA-seq activity predicted for the ALT sequence with the disrupted promoters. (c) Comparison of scores for REF sequence between RNA-seq and CAGE. The dashed line represents a linear fit. (d) Comparison of the ratio of scores (ALT/REF) between RNA-seq and CAGE, with color representing the size of the gene. Note, not the entire gene body was scored for longer genes and therefore, the indicated gene sizes reflect the width of the scored regions. (a-d) Each point represents one of the 2111 promoters analysed and the values are averages across the 20 tissues for which the scores were predicted. (e) Violin plots of ratio of scores (ALT/REF) at the TSS (sum of predicted CAGE signals in the CAGE scoring region) and at the exons (mean of predicted RNA-seq signal). Distal exons are binned for brevity. (f) Same as in e, except that exons are classified to those that overlap or lie downstream of the promoter region (Figure 1b). (g) Ratio of RNA-seq score (ALT/REF) only in exons downstream of the promoter region. Distances measured center-to-center from the last disrupted exon. (a-f) The number of exons representing the distributions are indicated. Note, y-axes are in log scale. (e-g) Very long genes with exons beyond the prediction region were not plotted.

To further explore this divergence between CAGE and RNA-seq track predictions, we focused on the GC-adjusted random base substitution approach. We also averaged the signals across the 20 scored tissues to avoid potential biases of tissue-specific expression or exon usage (Figure 2a-c). Firstly we confirmed that responding genes were not enriched in genes with highest expression in the reference scenario (Figure 2b). Given that the predicted CAGE and RNA-seq scores are linearly correlated for reference sequences as expected, we used the change in scores upon promoter disruption as a means to further investigate the divergence. We found an enrichment of longer genes amongst the non-responders (Figure 2d). This observation is in line with the previous reports of underestimation at longer distances, since distal exons of longer genes are much more separated from the disrupted promoter. To verify this, we scored the RNA-seq predictions separately for each exon, which revealed that the predicted effect size diminished rapidly with the exon number (Figure 2e). However, visual analysis of the predicted RNA-seq tracks of several non-responder genes revealed that all exons downstream of the disrupted region have similar magnitude of predicted RNA-seq expression, regardless of the exon rank (data not shown). Thus, segregating the scores into exons that are within the promoter region and those downstream (sequence unaffected) of it revealed a striking model behaviour (Figure 2f). The divergence between CAGE and RNA-seq predictions already appears to occur as early as at the second exon, if its sequence was not directly disrupted. In other words, it appears that whilst AlphaGenome recognises the loss of promoter elements in the form of reduced CAGE and RNA-seq signals at exons located within the span of disrupted promoter elements, it does not pay sufficient attention to the promoter while predicting the expression at downstream exons. Further analysis specific to downstream exons revealed that effect sizes are reduced at exons further away from the disruption (Figure 2g), which indicates that the model’s attention between promoters and exons is reduced at increasing distances, which is comparable to the observations of reduced effect sizes from distal enhancers^20,21^.

While these divergent predictions are in most circumstances at odds with transcriptional biology, there are foreseeable exceptions where they may indeed reflect biology. Firstly, some of the genes included in the analysis (due to random selection) are members of clustered gene families, which vary only at their first exon but share downstream exons (e.g., *PCDHA1-13*). In this scenario, it is biologically plausible that the disruption of the promoter of, say, *PCDHA3* does not abolish the (measured) expression at downstream exons, since they may be initiated by the other members of the gene family. Secondly, many genes feature alternate transcription start sites. While known transcription start sites are often associated with robust CAGE peaks and, therefore, disrupted in this study, visual examination of experimental CAGE tracks at many genes revealed weaker signals distributed throughout the gene body, which may indicate the presence of weak transcriptional initiation sites. It is possible that the residual RNA-seq signal predicted at the downstream exons is a result of transcriptional initiation from these sites. However, one would expect that this would also result in an associated increase in predicted CAGE signals at these weak transcription start sites, which we do not observe (e.g., Figure 1e between exons 3 and 4 of *CBX2*). Thirdly, random sequences have been reported to show promoter activity^35^, which may partly explain the residual activity (e.g., ALT/REF ratio in several cases > 0.1 for CAGE). However, this does not explain the uncoupling of correlation between CAGE and RNA-seq signals.

Several strategies could help address the limitations of current models in recapitulating true biology, specifically, in preserving the correlation between CAGE and RNA-seq; and in understanding the directionality/continuity of gene transcription. The most direct way, although currently difficult to implement at scale, would be to train S2F models on individual-specific genome sequences paired with matched functional genomics data. This could both alleviate the mismatch between the functional genomics data (usually obtained from cancer cell lines and patient samples) and the reference genome sequences used during training; as well as expose the models to naturally occurring genetic variation, including promoter variants and larger structural variants. Training data could also be augmented with experimentally informed promoter perturbations, or by model distillation which can improve variant-effect prediction^36^. A complementary approach would be to modify the training objective, for example by penalizing variation in predicted gene expression across exons of the same transcript, with the caveat of imposing assumptions that may not always reflect genuine biology, particularly in the presence of alternative promoter usage, transcript-specific regulation, or tissue-dependent isoform expression.

In summary, our findings show that S2F models do not necessarily recapitulate fundamental features of transcriptional biology, even when generating apparently plausible molecular predictions. Model outputs should therefore not be interpreted as biological evidence without appropriate validation. This is particularly important in diagnostic settings, in which inaccurate predictions could lead to the incorrect prioritization or classification of noncoding variants. Experimental validation and systematic benchmarking against established biological principles will remain essential for the responsible application of S2F models in human genetics.

## Data and code availability

The scripts used to perform the evaluations are available at https://github.com/SpielmannLab/S2F_PromoterDisruption. Note that the randomness in selection of genes analysed and the disruption of promoters may result in model predictions that deviate slightly from presented results. However, the general trends have remained consistent across multiple independent trials.

## Acknowledgements

M.S. is a DZKJ principal investigator and is supported by grants from the Deutsche Forschungsgemeinschaft (DFG) (DFG -SP 1532/3-2, DFG - SP 1532/13-1, DFG - SP 1532/16- 1, DFG - SP 1532/18-1, DFG - 515637292), the CRC 1665, the Einstein Foundation and the Max Planck Society.

## References

1. Sreenivasan, V.K.A., Yumiceba, V., and Spielmann, M. (2025). Structural variants in the 3D genome as drivers of disease. Nat. Rev. Genet. 10.1038/s41576-025-00862-x.

2. Balachandran, S., Prada-Medina, C.A., Mensah, M.A., Glaser, J., Kakar, N., Nagel, I., Pozojevic, J., Audain, E., Hitz, M.-P., Kircher, M., et al. (2024). STIGMA: Single-cell tissue-specific gene prioritization using machine learning. Am. J. Hum. Genet. 111, 338–349. 10.1016/j.ajhg.2023.12.011.

3. David, A., and Sternberg, M.J.E. (2023). Protein structure-based evaluation of missense variants: Resources, challenges and future directions. Curr. Opin. Struct. Biol. 80, 102600. 10.1016/j.sbi.2023.102600.

4. Adzhubei, I., Jordan, D.M., and Sunyaev, S.R. (2013). Predicting Functional Effect of Human Missense Mutations Using PolyPhenlJ2. Curr. Protoc. Hum. Genet. 76. 10.1002/0471142905.hg0720s76.

5. Siepel, A., Bejerano, G., Pedersen, J.S., Hinrichs, A.S., Hou, M., Rosenbloom, K., Clawson, H., Spieth, J., Hillier, L.W., Richards, S., et al. (2005). Evolutionarily conserved elements in vertebrate, insect, worm, and yeast genomes. Genome Res. 15, 1034–1050. 10.1101/gr.3715005.

6. Siepel, A., Pollard, K.S., and Haussler, D. (2006). New Methods for Detecting Lineage-Specific Selection. In Research in Computational Molecular Biology Lecture Notes in Computer Science., A. Apostolico, C. Guerra, S. Istrail, P. A. Pevzner, and M. Waterman, eds. (Springer Berlin Heidelberg), pp. 190–205. 10.1007/11732990_17.

7. Kircher, M., Witten, D.M., Jain, P., O’Roak, B.J., Cooper, G.M., and Shendure, J. (2014). A general framework for estimating the relative pathogenicity of human genetic variants. Nat. Genet. 46, 310–315. 10.1038/ng.2892.

8. Avsec, Ž., Agarwal, V., Visentin, D., Ledsam, J.R., Grabska-Barwinska, A., Taylor, K.R., Assael, Y., Jumper, J., Kohli, P., and Kelley, D.R. (2021). Effective gene expression prediction from sequence by integrating long-range interactions. Nat. Methods 18, 1196–1203. 10.1038/s41592-021-01252-x.

9. Linder, J., Srivastava, D., Yuan, H., Agarwal, V., and Kelley, D.R. (2025). Predicting RNA-seq coverage from DNA sequence as a unifying model of gene regulation. Nat. Genet. 57, 949–961. 10.1038/s41588-024-02053-6.

10. Pampari, A., Shcherbina, A., Kvon, E.Z., Kosicki, M., Nair, S., Kundu, S., Kathiria, A.S., Risca, V.I., Kuningas, K., Alasoo, K., et al. (2024). ChromBPNet: bias factorized, base-resolution deep learning models of chromatin accessibility reveal cis-regulatory sequence syntax, transcription factor footprints and regulatory variants. Preprint at Genetics, 10.1101/2024.12.25.630221 https://doi.org/10.1101/2024.12.25.630221.

11. De Almeida, B.P., Reiter, F., Pagani, M., and Stark, A. (2022). DeepSTARR predicts enhancer activity from DNA sequence and enables the de novo design of synthetic enhancers. Nat. Genet. 54, 613–624. 10.1038/s41588-022-01048-5.

12. Zhou, J. (2022). Sequence-based modeling of three-dimensional genome architecture from kilobase to chromosome scale. Nat. Genet. 54, 725–734. 10.1038/s41588-022-01065-4.

13. Avsec, Ž., Latysheva, N., Cheng, J., Novati, G., Taylor, K.R., Ward, T., Bycroft, C., Nicolaisen, L., Arvaniti, E., Pan, J., et al. (2026). Advancing regulatory variant effect prediction with AlphaGenome. Nature 649, 1206–1218. 10.1038/s41586-025-10014-0.

14. Shrikumar, A., Tian, K., Avsec, Ž., Shcherbina, A., Banerjee, A., Sharmin, M., Nair, S., and Kundaje, A. (2018). Technical Note on Transcription Factor Motif Discovery from Importance Scores (TF-MoDISco) version 0.5.6.5. Preprint at arXiv, 10.48550/ARXIV.1811.00416 https://doi.org/10.48550/ARXIV.1811.00416.

15. Alipanahi, B., Delong, A., Weirauch, M.T., and Frey, B.J. (2015). Predicting the sequence specificities of DNA- and RNA-binding proteins by deep learning. Nat. Biotechnol. 33, 831–838. 10.1038/nbt.3300.

16. Schreiber, J. (2026). Tangermeme: a toolkit for understanding cis-regulatory logic using deep learning models. Nat. Methods. 10.1038/s41592-026-03254-z.

17. Shrikumar, A., Greenside, P., and Kundaje, A. (2017). Learning Important Features Through Propagating Activation Differences. 10.48550/ARXIV.1704.02685.

18. Aksu, E.D., and Vingron, M. (2026). Context-aware sequence-to-function model of human gene regulation. Nat. Commun. 17, 6200. 10.1038/s41467-026-75527-2.

19. Cheng, J., Taylor, K.R., Nicolaisen, L., Pan, J., Bycroft, C., Perino, M., Ward, T., Hawkes, G., Covill, L.E., Weilert, M., et al. (2026). AlphaGenome Atlas: in silico mutagenesis of the entire human genome improves prioritization and interpretation of non-coding variants. Preprint at Genetic and Genomic Medicine, 10.64898/2026.09.16.26363192 https://doi.org/10.64898/2026.09.16.26363192.

20. Karollus, A., Mauermeier, T., and Gagneur, J. (2023). Current sequence-based models capture gene expression determinants in promoters but mostly ignore distal enhancers. Genome Biol. 24, 56. 10.1186/s13059-023-02899-9.

21. Murphy, A., and Koo, P.K. (2026). Benchmarking seq2func models on distal enhancer effects with CRISPRi screens. Genomics X AI Blog. 10.5281/zenodo.21383205.

22. Mostafavi, S., Tu, X., Spiro, A., and Chikina, M. (2026). A modality gap in personal-genome prediction by sequence-to-function models. Preprint at Bioinformatics, 10.64898/2026.02.01.702969 https://doi.org/10.64898/2026.02.01.702969.

23. Huang, D.-S., Li, B., Zhang, Q., and Bao, W. eds. (2027). GenBench: A Comprehensive Benchmark Dataset for Sequence-Based Genomic Models (Springer Nature Singapore) 10.1007/978-981-92-3498-1.

24. Huang, C., Shuai, R.W., Baokar, P., Chung, R., Rastogi, R., Kathail, P., and Ioannidis, N.M. (2023). Personal transcriptome variation is poorly explained by current genomic deep learning models. Nat. Genet. 55, 2056–2059. 10.1038/s41588-023-01574-w.

25. Sasse, A., Ng, B., Spiro, A.E., Tasaki, S., Bennett, D.A., Gaiteri, C., De Jager, P.L., Chikina, M., and Mostafavi, S. (2023). Benchmarking of deep neural networks for predicting personal gene expression from DNA sequence highlights shortcomings. Nat. Genet. 55, 2060–2064. 10.1038/s41588-023-01524-6.

26. Drusinsky, S., and Pollard, K.S. (2026). Causal variant underestimation is a major overlooked driver of sequence-to-function model underperformance. Preprint at Genetics, 10.64898/2026.09.08.750172 https://doi.org/10.64898/2026.09.08.750172.

27. Haberle, V., and Stark, A. (2018). Eukaryotic core promoters and the functional basis of transcription initiation. Nat. Rev. Mol. Cell Biol. 19, 621–637. 10.1038/s41580-018-0028-8.

28. Gilbert, L.A., Larson, M.H., Morsut, L., Liu, Z., Brar, G.A., Torres, S.E., Stern-Ginossar, N., Brandman, O., Whitehead, E.H., Doudna, J.A., et al. (2013). CRISPR-Mediated Modular RNA-Guided Regulation of Transcription in Eukaryotes. Cell 154, 442–451. 10.1016/j.cell.2013.06.044.

29. Juven-Gershon, T., and Kadonaga, J.T. (2010). Regulation of gene expression via the core promoter and the basal transcriptional machinery. Dev. Biol. 339, 225–229. 10.1016/j.ydbio.2009.08.009.

30. Weingarten-Gabbay, S., Nir, R., Lubliner, S., Sharon, E., Kalma, Y., Weinberger, A., and Segal, E. (2019). Systematic interrogation of human promoters. Genome Res. 29, 171–183. 10.1101/gr.236075.118.

31. Dynan, W.S., and Tjian, R. (1985). Control of eukaryotic messenger RNA synthesis by sequence-specific DNA-binding proteins. Nature 316, 774–778. 10.1038/316774a0.

32. FitzGerald, P.C., Shlyakhtenko, A., Mir, A.A., and Vinson, C. (2004). Clustering of DNA Sequences in Human Promoters. Genome Res. 14, 1562–1574. 10.1101/gr.1953904.

33. FANTOM Consortium and the RIKEN PMI and CLST (DGT), Forrest, A.R.R., Kawaji, H., Rehli, M., Baillie, J.K., de Hoon, M.J.L., Haberle, V., Lassmann, T., Kulakovskiy, I.V., Lizio, M., et al. (2014). A promoter-level mammalian expression atlas. Nature 507, 462–470. 10.1038/nature13182.

34. Jiang, M., Anderson, J., Gillespie, J., and Mayne, M. (2008). uShuffle: A useful tool for shuffling biological sequences while preserving the k-let counts. BMC Bioinformatics 9, 192. 10.1186/1471-2105-9-192.

35. De Boer, C.G., Vaishnav, E.D., Sadeh, R., Abeyta, E.L., Friedman, N., and Regev, A. (2020). Deciphering eukaryotic gene-regulatory logic with 100 million random promoters. Nat. Biotechnol. 38, 56–65. 10.1038/s41587-019-0315-8.

36. Wang, L., Galvin, J., Vaz, E., Keener, R., and Battle, A. (2026). Synthetic MPRA perturbation supervision improves variant-effect prediction in sequence-to-function models. Preprint at Genomics, 10.64898/2026.11.02.763699 https://doi.org/10.64898/2026.11.02.763699.

